# Diurnal profiles of physical activity and postures derived from wrist-worn accelerometry in UK adults

**DOI:** 10.1101/600650

**Authors:** Ignacio Perez-Pozuelo, Thomas White, Kate Westgate, Katrien Wijndaele, Nicholas J. Wareham, Soren Brage

**Affiliations:** MRC Epidemiology Unit, University of Cambridge, Cambridge, United Kingdom

**Author notes:** Address for correspondence: Dr. Søren Brage, MRC Epidemiology Unit University of Cambridge School of Clinical Medicine, Box 285, Institute of Metabolic Science Cambridge Biomedical Campus Cambridge CB2 0QQ, United Kingdom.

**Keywords:** Pitch angle, Roll angle, Movement, Signal Processing, BMI, PAEE, Sedentary Behaviour

## Abstract

**Background:** Wrist-worn accelerometry is the commonest objective method for measuring physical activity in large-scale epidemiological studies. Research-grade devices capture raw triaxial acceleration which, in addition to quantifying movement, facilitates assessment of orientation relative to gravity. No population-based study has yet described the interrelationship and variation of these features by time and personal characteristics.

**Methods:** 2043 UK adults (35-65years) wore an accelerometer on the non-dominant wrist and a chest-mounted combined heart-rate-and-movement sensor for 7days free-living. From raw (60Hz) wrist acceleration, we derived movement (non-gravity acceleration) and pitch and roll (arm) angles relative to gravity. We inferred physical activity energy expenditure (PAEE) from combined sensing and sedentary time from approximate horizontal arm-angle coupled with low movement.

**Results:** Movement differences by time-of-day and day-of-week were associated with arm-angles; more movement in downward arm-positions. Mean(SD) movement was similar between sexes ∼31(42)mg, despite higher PAEE in men, 53(22) vs 48(19)J·min^-1^·kg^-1^. Women spent longer with the arm pitched >0° (53% vs 36%) and less time at <0° (37% vs 53%). Diurnal pitch was 2.5-5° above and 0-7.5° below horizontal during night and daytime, respectively; corresponding roll angles were ∼0° and ∼20° (thumb-up). Differences were more pronounced in younger participants. All diurnal profiles indicated later wake-times on weekends. Daytime pitch was closer to horizontal on weekdays; roll was similar. Sedentary time was higher (17 vs 15hours/day) in obese vs normal-weight individuals.

**Conclusions:** More movement occurred in arm positions below horizontal, commensurate with activities including walking. Findings suggest time-specific population differences in behaviours by age, sex, and BMI.

## BACKGROUND

Wrist-worn accelerometry has become a feasible option for the objective measurement of physical activity in large-scale epidemiological studies, such as Pelotas birth cohorts, the UK Biobank and Whitehall II (1–3). Additionally, public adoption of consumer-grade wearable devices that include accelerometry has been increasing steadily in recent years (4–6), with potential utility for public health research (7).

Accelerometers record a continuous time-series of data and recent advances in the technology and battery life allow for ubiquitous capture of raw accelerometer signals which have the potential to provide insights to interventional and epidemiological studies. Several features can be easily extracted from the acceleration signal, including the magnitude of movement and the orientation of the accelerometer with respect to gravity.

Previous research using wrist accelerometry has described variation in population physical activity expressed predominantly as the activity-related acceleration magnitude. For example, da Silva et al noted age and sex differences in three Brazilian birth cohorts from Pelotas assessed at the ages 6, 18, and 30 years of age (1), and Doherty et al described the unique diurnal patterns of physical activity by age group, documenting that the lower activity levels generally observed in older adults are particularly pronounced in the later hours in the day (2). Magnitude-based measures of activity have also been related to health outcomes, such as body composition and fitness (7,8).

Less attention has been given to the description of orientation-related measures of human behaviour. Pitch and roll angles are examples of well-defined, biomechanically relevant and easy-to-interpret signal features that describe device orientation. In figure 1, we illustrate pitch and roll for an individual with an accelerometer placed on the left wrist, and axes aligned as shown. Body posture is by definition a description of angles of all segments of the body which can in theory all be measured but in practice usually are not in studies of free-living behaviour. However, as body segments are connected, and therefore range of motion is restricted, measurements and their derivatives are highly correlated (9). This allows inferences from the measurement of one body site to be made on whole-body posture. For example, previous work has shown strong correlations between time spent sedentary inferred from wrist accelerometry (by combining information on acceleration magnitude and pitch angle) and thigh accelerometry (r∼0.93) (10).

**Figure 1:**
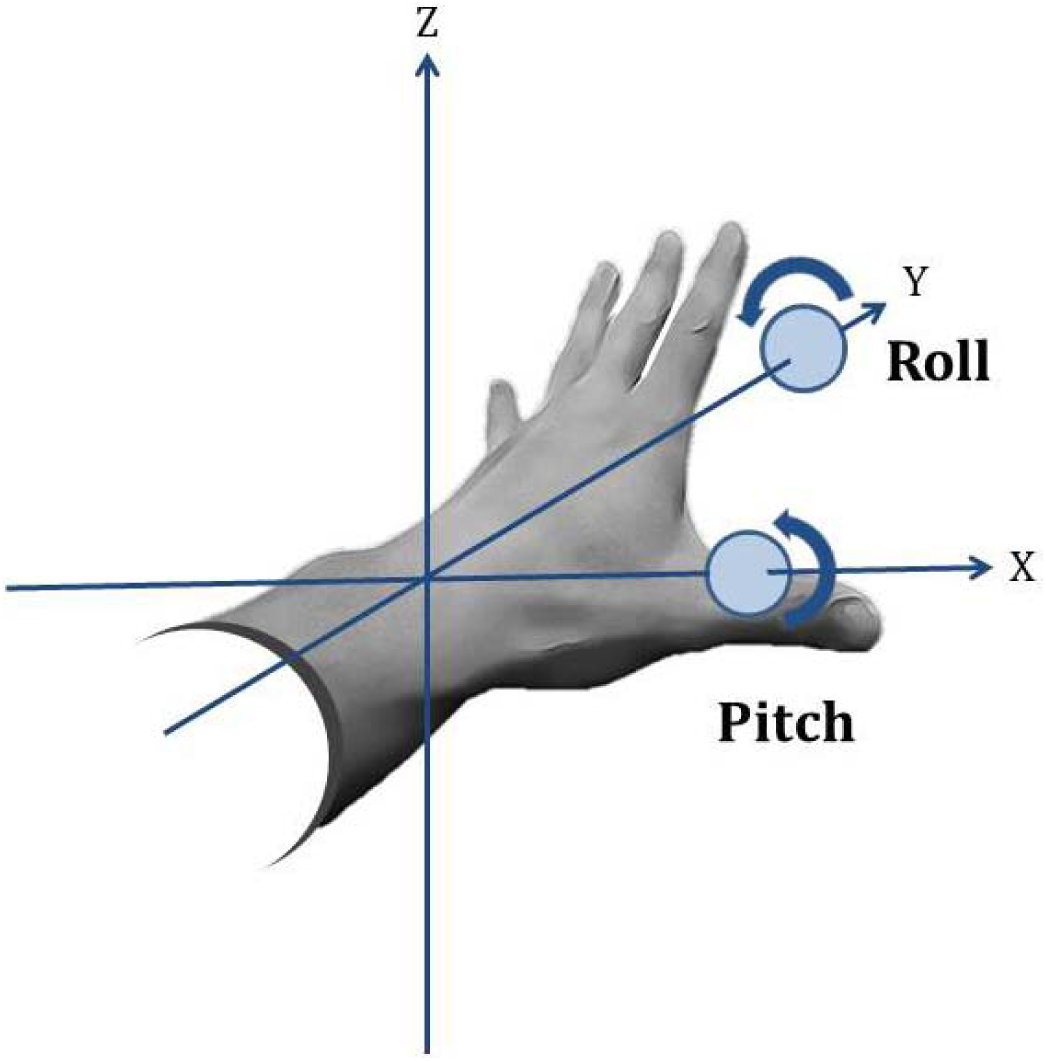
Schematic of Pitch and Roll on participant with accelerometer on the left wrist, including axes alignment. Roll is defined by rotation around the Y axis, while Pitch is defined by rotation around the X axis. (Note that axis labelling depends on study protocol and device specifications).

Sedentary behaviour can be defined as any waking behaviour that is characterized by an energy expenditure ≤ 1.5 METs while the subject is engaging in either sitting, lying or reclining postures (11). People spend the majority of their time in sedentary behaviours, and the proportion of time spent sedentary increases as people age (12). High volumes of sedentary behaviour have been associated with increased mortality and risk of developing chronic conditions (12–16). This only seems to be eliminated by very high levels of moderate intensity physical activity (60-75 min per day, i.e. equivalent to double the amount currently recommended for adults (14)). However, most of this evidence base is based on self-reported sedentary and activity estimates which come with important methodological limitations and bias (17).

Consequently, objectively assessing sedentary behaviours, as well as characterizing different activities performed during daily living may be critical to inform public health recommendations. Traditionally, sedentary and active behaviours were characterized using such intensity derived measures from the accelerometer signal. Supplementary Figure 1 provides a visual representation of triaxial wrist acceleration (top panel) during four common activities of lying, walking, sitting, and cycling, alongside derived pitch and roll angles (bottom panel), demonstrating clear differences between activity types. When assessing activity patterns, diurnal profiles of pitch and roll combined with movement intensity metrics may allow us to further understand how different postures relate to different activities and activity intensities.

In this study, we describe the distribution of wrist postures, acceleration, derived sedentary time and PAEE in a large cohort of UK adults (n=2043 participants). These analyses allow us to further understand the distribution of sedentary and active behaviours in the population and how this distribution may differ based on time of the day, sex, age, body mass index (BMI) and some other substrata. Ultimately, the methodology developed for the work presented aims to help inform how changes in sedentary and active behaviours may impact energy expenditure.

## METHODS

### Study Population

The Fenland Study is an ongoing prospective cohort study of 12,435 men and women aged 35-65 years, designed to identify the behavioural, environmental and genetic causes of obesity and type-2 diabetes. As previously described in detail, participants attended one of three clinical research facilities in the region surrounding Cambridge, UK, and completed a series of physical assessments and questionnaires (18). Exclusion criteria for participation in the study were: clinically diagnosed diabetes mellitus, inability to walk unaided, terminal illness, clinically diagnosed psychotic disorder, pregnancy or lactation. Following the baseline clinic visit, all participants were asked to wear a combined heart rate and movement sensor (Actiheart, CamNtech, Cambridgeshire, UK) for 6 consecutive days and nights, and a subsample of 2100 participants were asked to simultaneously wear a wrist accelerometer (GeneActiv, ActivInsights, Cambridgeshire, UK) on the non-dominant wrist. This subsample constitutes the sampling frame for the current analyses. Participants were excluded from this analysis if they had insufficient individual calibration data, or had less than 72h of concurrent wear data (equivalent of 3 full days of recording). Given only very few participants were very severely underweight (BMI ≤ 15) in this subset of the Fenland study, they were also excluded, resulting in a total of 2043 subjects.

All participants provided written informed consent and the study was approved by the local research ethics committee (NRES Committee – East of England Cambridge Central) and performed in accordance with the Declaration of Helsinki.

### Data Collection

#### Physical activity measures

The combined heart rate and movement sensor attached to the participant’s chest, measured heart rate and uniaxial acceleration of the trunk in 15-second intervals (19). The wrist accelerometer worn on the non-dominant wrist recorded triaxial acceleration at 60 Hertz. Participants were instructed to wear both waterproof monitors continuously for 6 full days and nights during free-living conditions, including during showering and while they were sleeping.

During the baseline clinic visit, participants performed a ramped treadmill test to establish their individual heart rate response to a submaximal exercise test (20). These measurements produced calibration parameters that were used in a branched equation model of PAEE (21). Heart rate data collected during free-living was pre-processed to eliminate potential noise (22), following which the branched equation model was applied to calculate instantaneous PAEE (J·min^-1^·kg^-1^). This inference has been validated against intensity from indirect calorimetry (23,24) and volume from doubly-labelled water in several populations (25), including a sample of UK men and women in whom the technique was shown to explain 41% of the variance in free-living PAEE as well as no mean bias (26).

The wrist accelerometer data was processed using pampro, an open-source software package (27). The triaxial acceleration was auto-calibrated to local gravitational acceleration using a method described elsewhere (28). Non-wear time was defined as time periods where the standard deviation of the acceleration in each of the three axes fell below 13mg for over an hour, inferring that the device was completely stationary (29). When a non-wear period was detected, it was removed from the analyses. The magnitude of acceleration was calculated using *Vector Magnitude* (VM) (expressed in milli-g/mg) per sample:

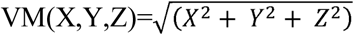

VM, or Euclidean Norm, can be interpreted as the magnitude of acceleration the device was subjected to at each measurement, which includes gravitational acceleration. Any potential noise component in the high-frequency domain was filtered out by a 20 Hertz low-pass filter. To isolate the movement-related acceleration, we also applied a high-pass Butterworth filter to the VM signal at 0.2 Hertz (therefore treating gravity as a low-frequency component) naming the resulting metric Vector Magnitude High-Pass Filtered (VM HPF, expressed in mg)(7,29). VM HPF is commonly used as a proxy of acceleration resulting from human movement, has high validity (30), and was the primary description of wrist movement in the following analyses.

When movement-related acceleration is removed by a low-pass filter (0.2 Hertz) to each of the three axes (X, Y and Z), the residual acceleration signal can be interpreted as a measurement of the rotated gravitational field vector which can then be used to determine the accelerometer’s pitch and roll orientation angles. Pitch and roll of the device were derived according to these formulae:

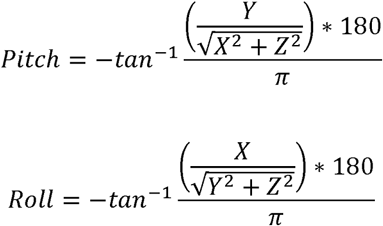

As the monitor was mounted in such a way that the X-axis was aligned in anatomically opposite directions for left-and right-handed participants, we multiplied it by −1 for all left-handed participants who wore the monitor on their right wrists to align with the anatomical coordinate system defined above (examples of untransformed data shown in supplementary figure 2). Consequently, positive pitch indicates upwards position of the arm (hand above elbow), while positive roll indicates the lateral (radial, thumb) side of the arm being higher than the medial (ulnar, pinky) side of the arm.

All derived signals were summarized to a common time resolution of one observation per hour. This window length was chosen since we were mostly interested in observing changes at a diurnal level, rather than variations within the hour.

Using the combined-sensing measurements, participants were stratified by average activity energy expenditure: lower active (≤ 39 J · min^-1^ · kg^-1^), medium (40-56 J · min^-1^ · kg^-1^) and upper (>=57 J · min^-1^ · kg^-1^). These activity estimates were calculated for each participant for each day of the week and then averaged, allowing us to generate a picture of changes in behaviour over the course of the week.

Similarly, we calculated estimates of time spent in sedentary (i.e. sitting or reclining) by detecting bouts where wrist pitch (i.e. arm elevation) is ≥ 15 ° below the horizontal, while wrist acceleration is minimal (VM HPF ≤ 47.61 mg)). This is based on principles from previously developed methodology which derives sedentary time estimates from wrist accelerometry data (i.e. sedentary sphere methodology (10)), as well as estimations of physical activity energy expenditure in free-living using wrist accelerometry (7). The latter defined the acceleration threshold (VM HPF =47.61 mg) equivalent to 1.5 gross METs (PAEE=35.5J.min^-1^.kg^-1^) as the cut-off for sedentary behaviour (7). Data in lower latitudes, that is, less than −15° from the horizontal, suggest hanging of the arm, associated to standing behaviours and are hence not classified as sedentary time. Equally, if the mean levels (VM HPF) over a minute fell into the light, moderate or vigorous category, they were not classified as sedentary behaviour.

Using the diurnal profiles derived from the cohort, we studied differences based on sex, age, activity levels, BMI and time of the day.

### Statistical analyses

We computed descriptive statistics (mean, median, standard deviation, minimum, maximum and variance) for the participants in this analysis. We examined wear-time distributions using the Friedman test for time-of-day (00:00-05:59, 06:00-11:59, and so on in six-hour periods) and tested the differences in weekdays versus weekend days using Wilcoxon signed ranks. These tests were performed in men and women separately. Mean acceleration differences (VM HPF) were examined using ANOVA for time of the day and day of the week. Differences between men and women are shown by using box plots, providing information about the median, inter-quartile range, minimum and maximum. We analysed the differences between different BMI groups (underweight ≤ 18.5 kg/m^2^, normal weight 18.5-24.9 kg/m^2^, overweight 25-29.9 kg/m^2^, obese 30-34.9 kg/m^2^ and severely obese ≥ 35 kg/m^2^) in both sexes based on pitch, roll, VM HPF and PAEE. Similarly, we conducted the analysis based on age group and PAEE levels. These summary statistics were computed at an hourly level after collapsing information derived on a fifteen-second time window.

Furthermore, we tested for differences in time spent in sedentary time across the different BMI populations using 3-way ANOVA and adjusting for age and sex.

Statistical tests were performed using Python (3.6.2) and Stata (v14, StataCorp, TX, USA).

## RESULTS

Among the 2043 participants, a total of 286,020 person-hours were included in our analysis, or an average of 5.8 days per participant. As shown in Table 1, PAEE was higher in men although both groups had large standard deviations. However, wrist movement was similar between genders but mean BMI was larger in men than in women for this cohort.

**Table 1:**
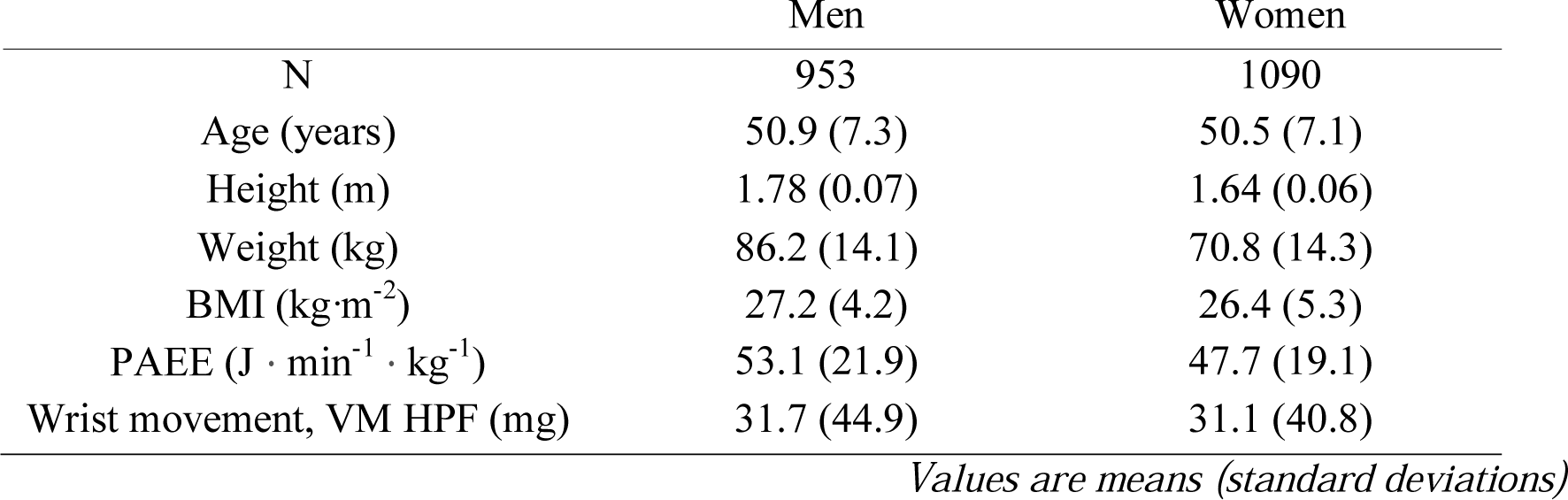
Characteristics of participants by sex (n=2043)

|  | Men | Women |
| --- | --- | --- |
| N | 953 | 1090 |
| Age (years) | 50.9 (7.3) | 50.5 (7.1) |
| Height (m) | 1.78 (0.07) | 1.64 (0.06) |
| Weight (kg) | 86.2 (14.1) | 70.8 (14.3) |
| BMI (kg·m<sup>-2</sup>) | 27.2 (4.2) | 26.4 (5.3) |
| PAEE (J · min<sup>-1</sup> · kg<sup>-1</sup>) | 53.1 (21.9) | 47.7 (19.1) |
| Wrist movement, VM HPF (mg) | 31.7 (44.9) | 31.1 (40.8) |

*Values are means (standard deviations)*

Figure 2A shows pitch and roll distributions for men and women; a 2-dimensional plot of pitch and roll is shown in supplementary figure 3. There is higher occurrence of pitch and roll positions around 0° and the roll distribution is distinctly bimodal with an additional peak around 35°. Less common are extreme anatomical wrist positions e.g., arms up in the air, reflected by a pitch >60°, or the radial (thumb) side of the arm turned inwards and downwards as indicated by less roll data below −45°. Figure 2B and 2C shows the differences among different age groups for average sedentary time and PAEE respectively. PAEE declines with age in both men and women, and there is a tendency for the wrist measure of sedentary time to increase with age, showing a close inverse relationship between these two measures.

**Figure 2:**
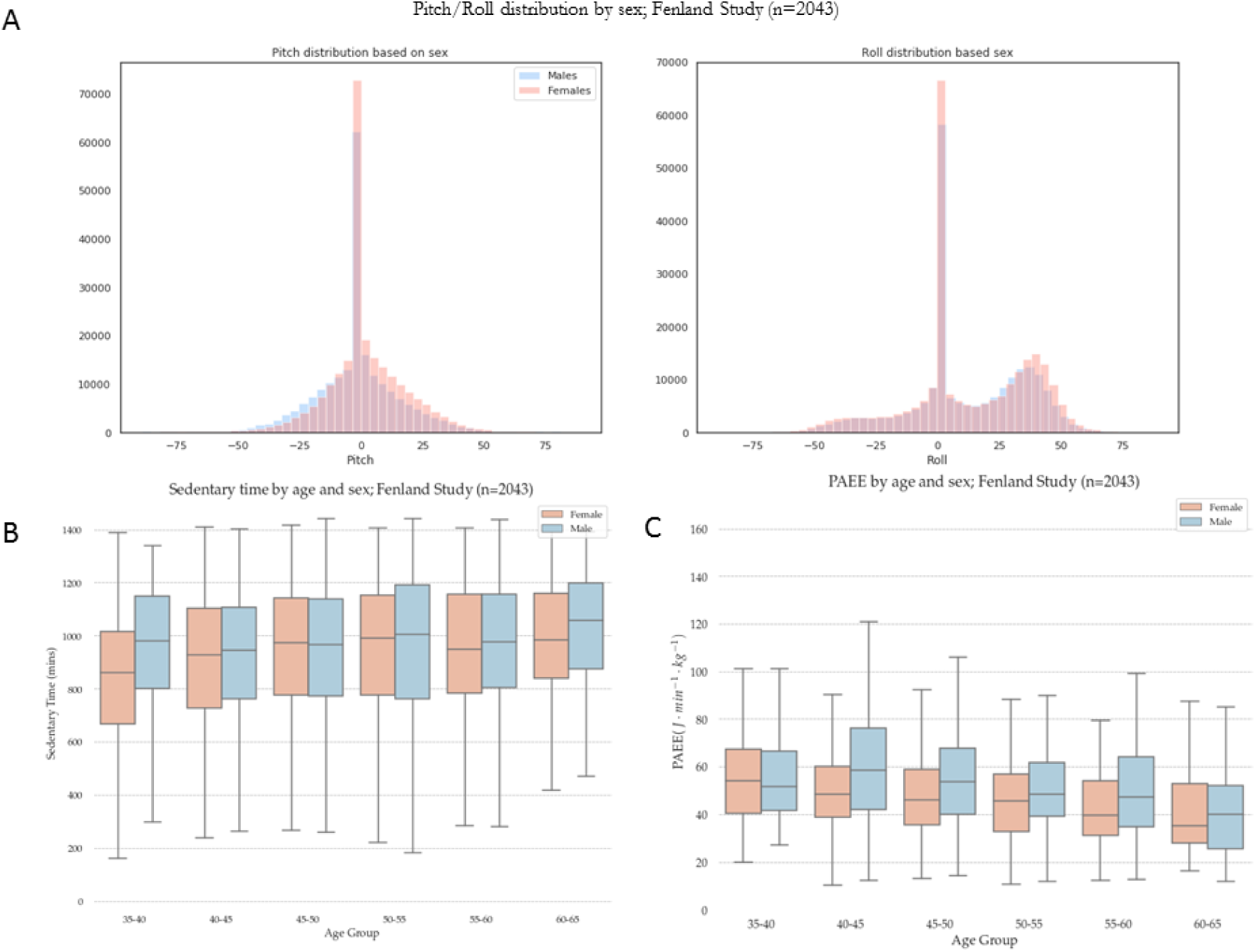
Pitch and roll (A) distribution among participants, and box plots for time spent sedentary (B) and PAEE (C) by age group and sex (n=2043).

### Relation between wrist movement and postures, and physical activity energy expenditure

Figure 3 shows differences in wrist measures by tertile of physical activity energy expenditure; more active individuals spend more time in low-pitch (below horizontal) postures; less active participants tend to be spending more time in postures that suggest sedentary behaviours, such as sitting or reclining. Whilst roll angles differ by activity level in women, there is almost no difference between groups in men; differences in wrist movement, however, are very clear in both genders.

**Figure 3:**
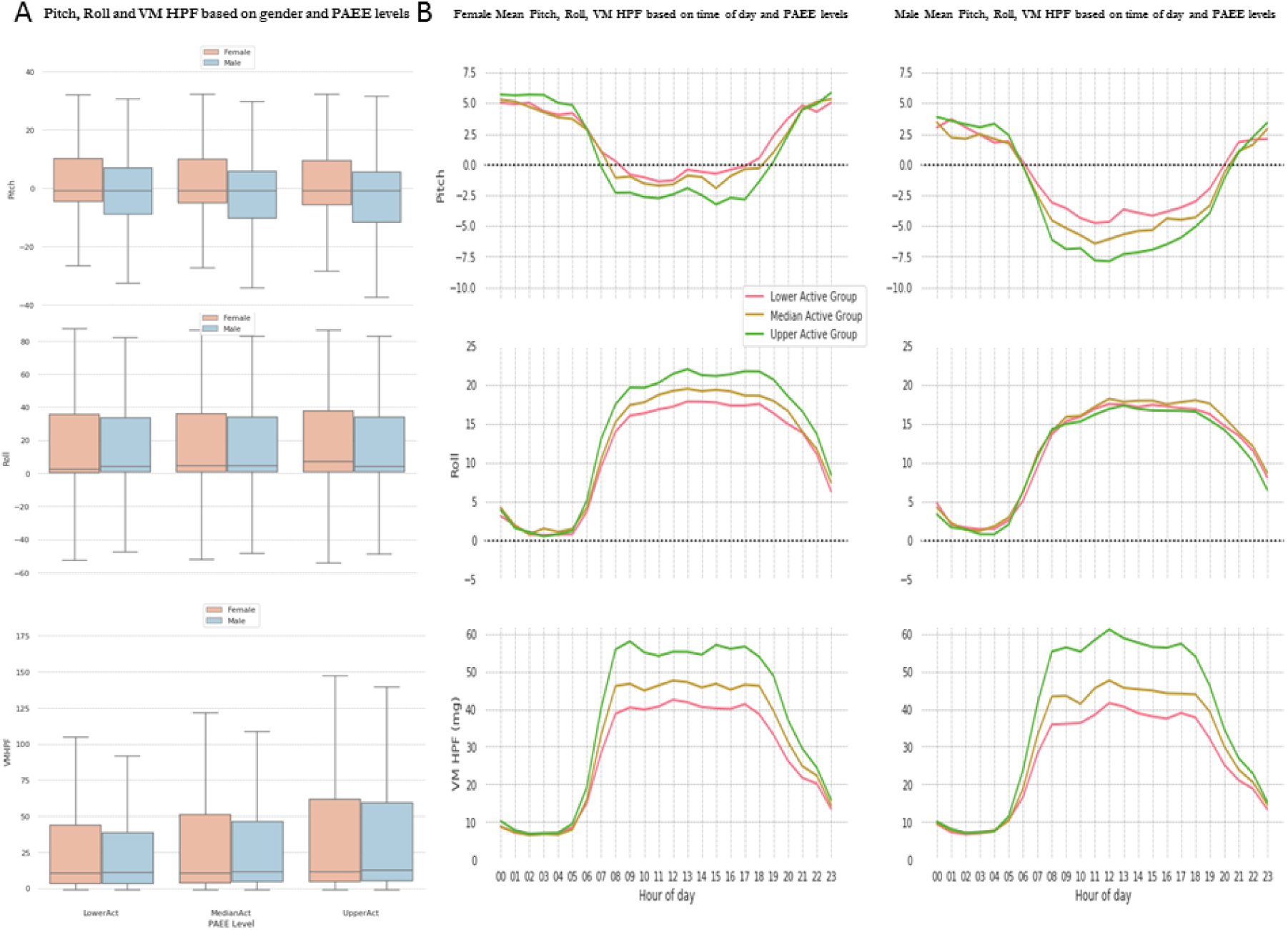
Pitch (top panels), Roll (middle panels) and Vector Magnitude High-Pass Filtered (VM HPF) by PAEE level (lower, medium or upper) and gender (A), and diurnal profiles by time of day in women and men (B).

Some of the most visually striking results regarding the role of posture on physical activity behaviours can be seen in the 3-dimensional time-lapse plots that appear on the online supplementary online material of this paper (see video). A schematic representation of these time-lapses is presented in figure 4 at four times of the day.

**Figure 4:**
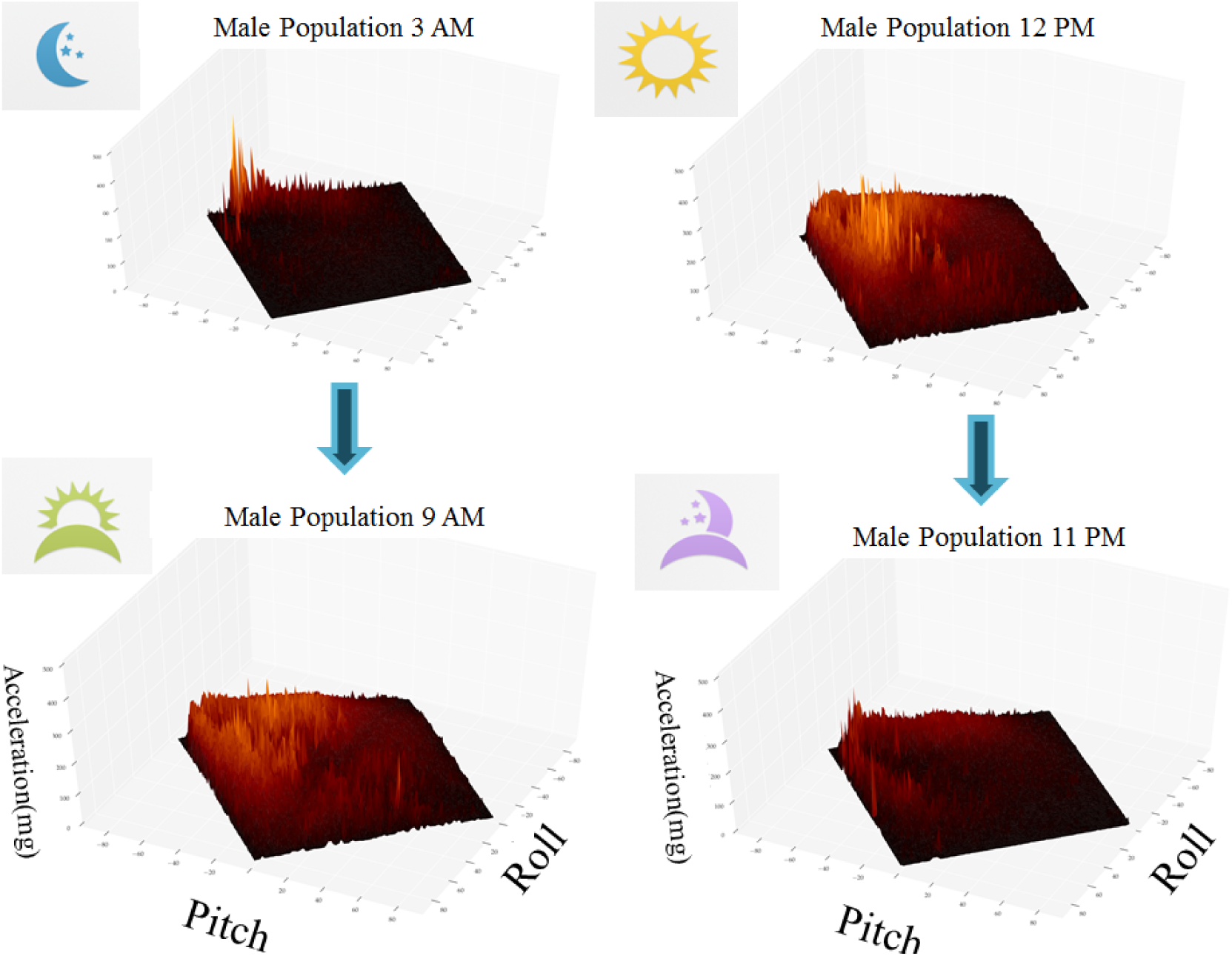
Schematic representation of time-lapse diurnal change in Pitch and Roll angular profiles and their associated acceleration signal (VM HPF, in mg). All plots have been normalized. (Figure derived from the male population of this analysis n=953). Full videos for both genders available in online supplemental material.

### Diurnal Profile Differences by sex and age

Figure 5 shows the distribution of pitch, roll, and movement intensity across the day, stratified by sex and age group. We observe differences between age groups within sex, but also differences between men and women within age groups. Most differences between men and women occur during the working hours (8 AM to 6PM) of the day, with little differences at night although women generally keep their arms at slightly higher pitch throughout the 24 hours. Some of the biggest differences between age groups in both sexes happen during the early hours of the morning and late hours of the evening. Arm angles differ more between age groups in men (lower pitch in older during working hours), and gender differences in pitch and roll profiles are most apparent among the 35-40 age group.

**Figure 5:**
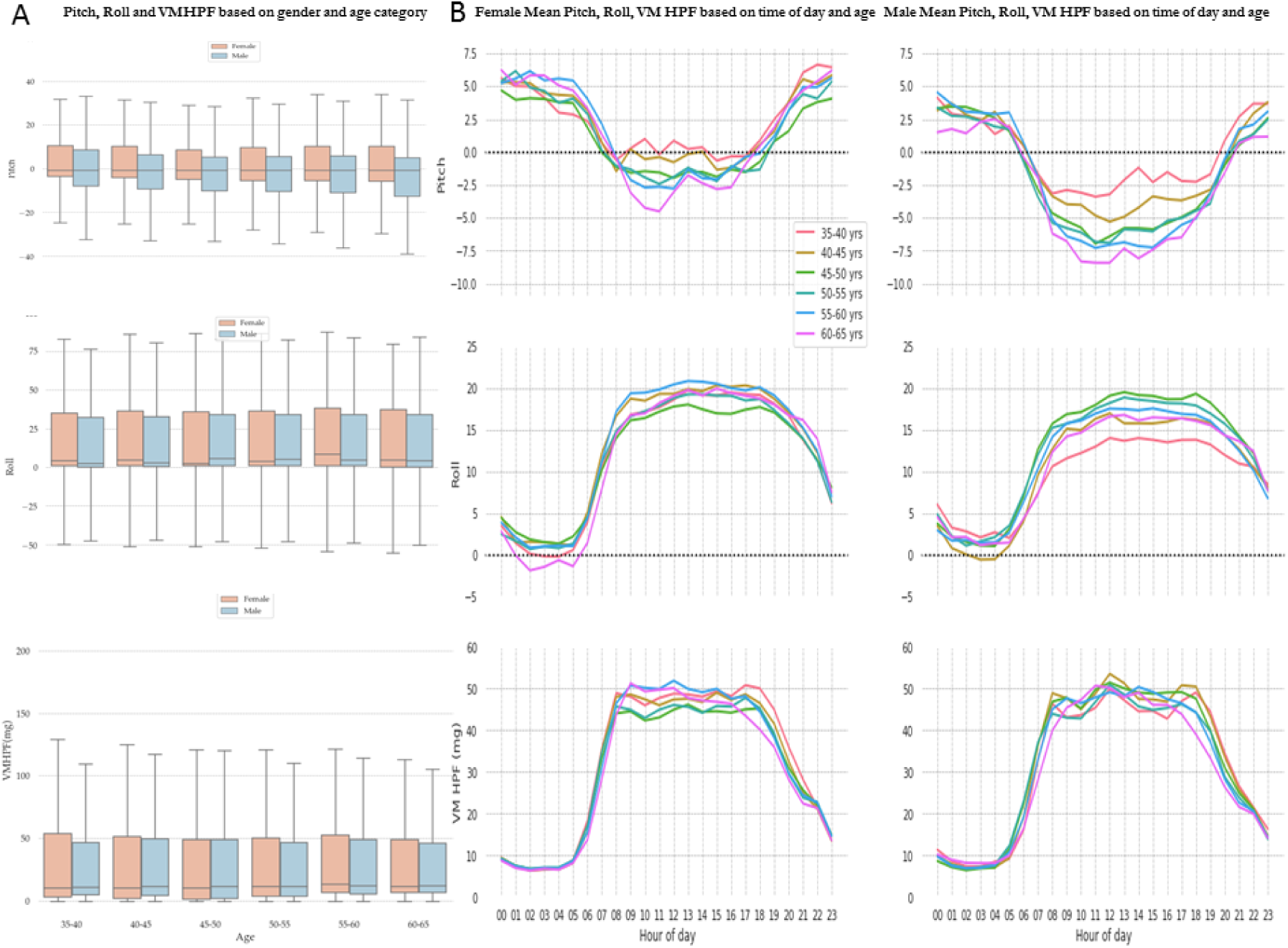
Pitch (top panels), Roll (mid panels) and VM HPF (bottom panels) profiles by time of the day and age group (from 35-40 to 60-65 years old) in women (middle column) and men (right column). Left column (A) shows participant-level summary data.

### Pitch and Roll Profiles Differ on Weekends versus Weekdays

Figure 6 shows average pitch, roll and movement intensity across the day, separately for each day of the week, and stratified by sex. The variation between weekdays at a population level is minimal, but they differ from the diurnal profiles at the weekend and particularly among sexes. A visible shift on weekend days towards later hours of the morning suggests a “*later start”* to the day, and later bed times on Friday and Saturday nights. The most extreme postural contrast are seen for pitch angles in men which reach the lowest level at the weekend (around −10°) in parallel to highest level of movement; pitch in women is also lower in the weekend but only to the weekday level of the men (around −5°) but with a similar level of movement as men.

**Figure 6:**
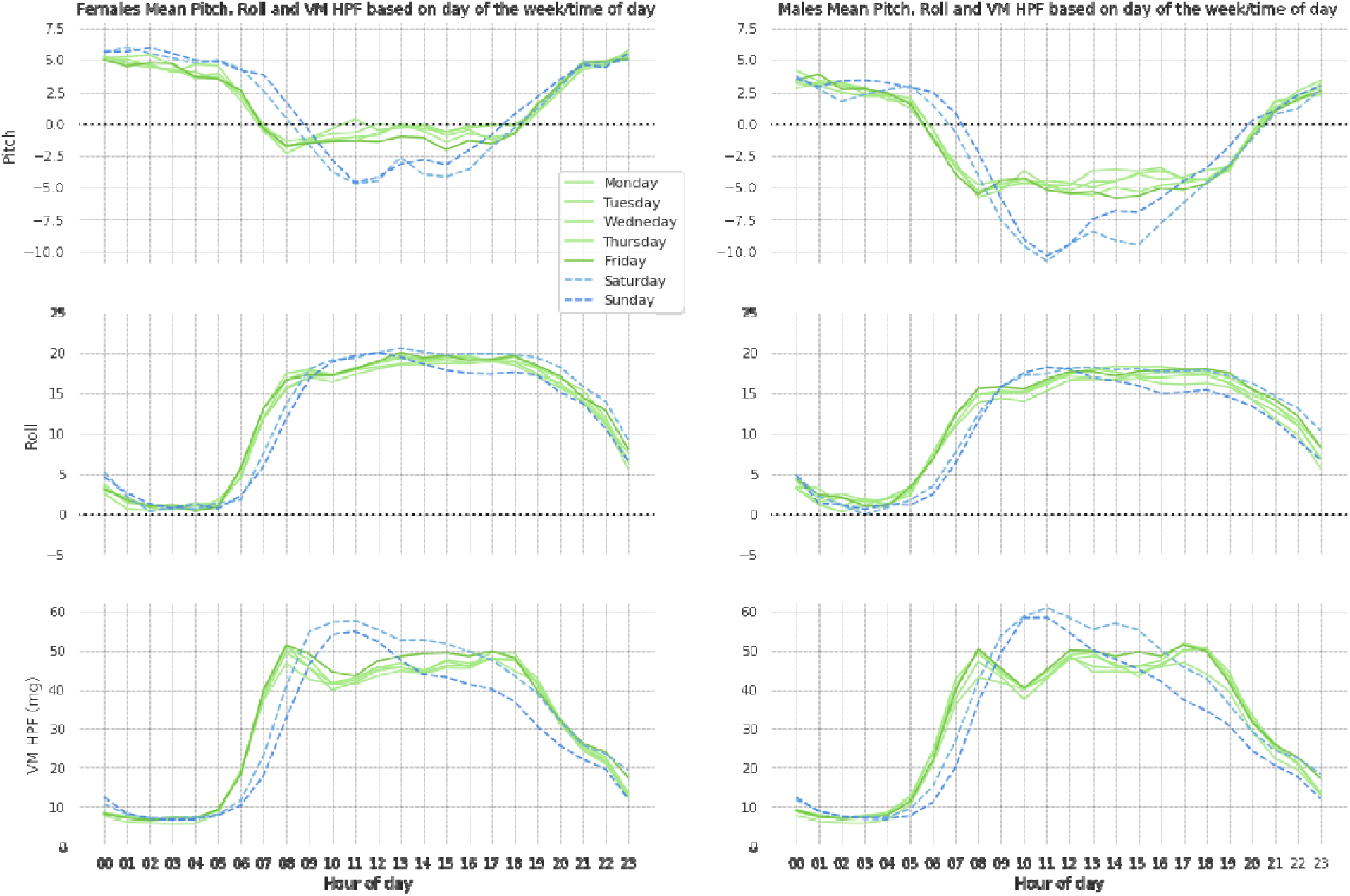
Differences in Pitch, Roll and VM HPF based on day of the week (weekdays in green, weekends in blue) and time of the day in women and men.

### Wrist Accelerometry Profiles by Gender and BMI

The differences in mean VM HPF between the different BMI groups are striking with obese individuals moving considerably less than normal-weight but equally notable are differences in pitch and roll profiles (Figure 7). Differences among groups were more apparent in men than in women when considering the diurnal profile. Somewhat surprisingly, given higher movement is generally occurring at the lower pitch angles (figure 3), overweight and obese individuals spend more time with their arms in this space but they just do not seem to move as much. The underweight women’s pitch and roll profile are very different to that observed in the severely obese men, suggesting that the higher level of mean physical activity in this group is also related to a very different set of activities. These observations are supported by stark differences on the average time spent in sedentary behaviours stratified by sex and BMI category, where non-obese participants spent considerably less time in sedentary behaviours than obese participants, particularly women. Also, the profiles observed in obese men closely resemble that observed in the oldest age group as presented in figure 5. We confirmed differences across different BMI groups for average time spent in sedentary behaviours, following adjustment for age and sex. We found that moderately obese participants spent significantly more time in sedentary behaviours than normal-weight participants (p<0.001), and so did severely obese (p<0.001) and even overweight participants (p<0.001). We also found a strong significant difference between overweight and moderately obese participants (p=0.0001); however, differences between normal-weight and underweight participants were not statistically significant (p=0.57).

**Figure 7:**
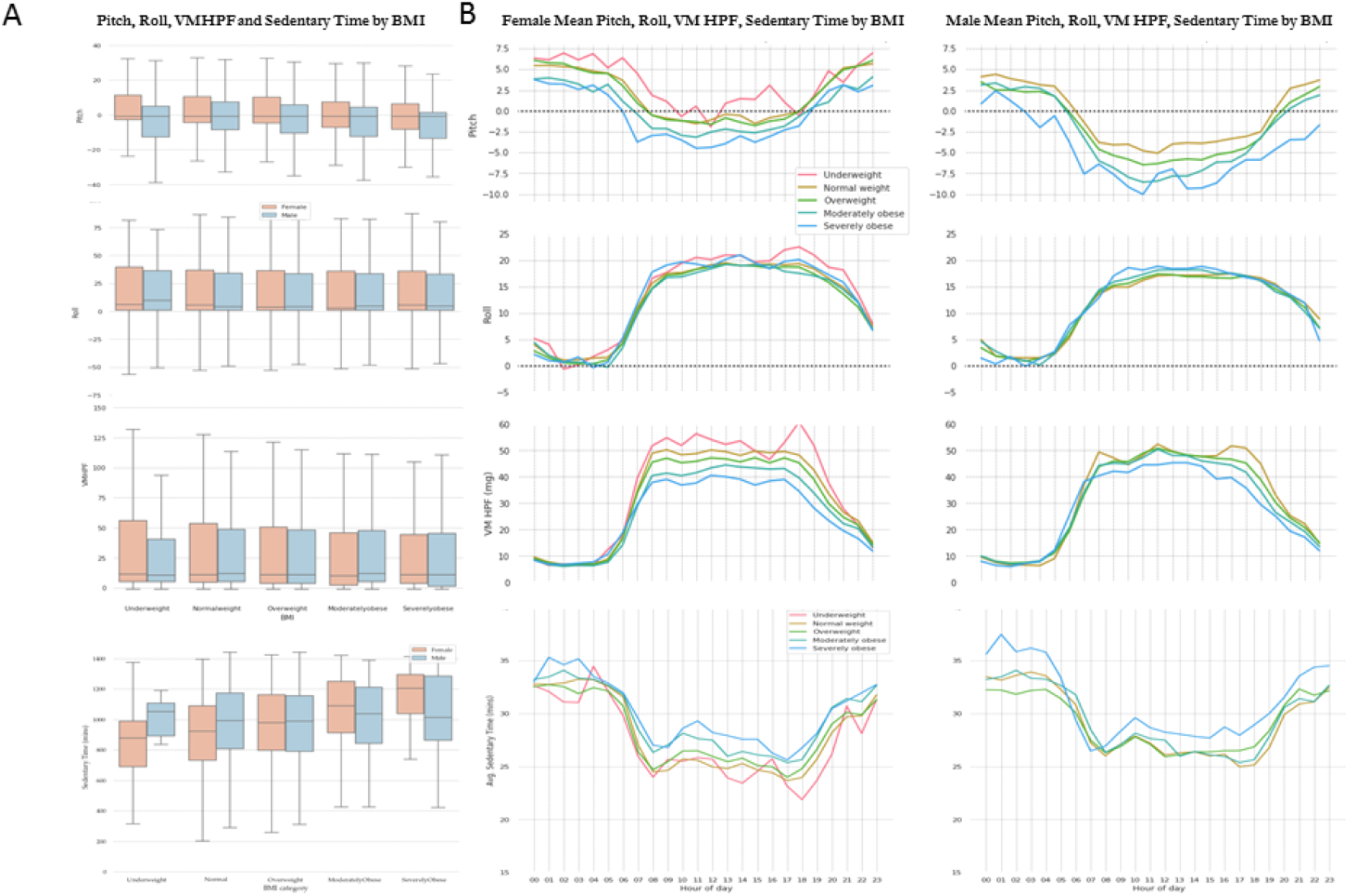
Pitch (top panels), Roll (second row panels), VM HPF (third row panels), and sedentary time (bottom row panels) profiles by time of the day in women and men, stratified by BMI categories (ranging from *underweight (BMI: 16-18.5)* to *severely obese (BMI*≥3*5)*).

## DISCUSSION

In this paper, we have explored the physical space in which physical activity occurs and described population differences in wrist movement and posture between men and women, age groups, BMI categories, and physical activity levels in a population sample of UK adults. Although higher activity was associated with lower pitch profiles, we observed the apparent paradox that older and more obese individuals who as groups are generally less active also spend more time at these postures, indicating that these groups either perform different types of activities or perform them at slower pace.

Vector magnitude of movement intensity and pitch-roll angular features can all be considered direct measures of human behaviour, rather than estimates, as there is very little inference involved in deriving them; they have biomechanical meaning in their own right as also illustrated in supplementary figure 1. The estimate of sedentary time, on the other hand, is not a direct measure but an estimate resulting from an inference but we have included it here to demonstrate the utility of combining directly measured features. Including movement as well as pitch, roll (both indicating posture), and sedentary time estimates in our analysis allowed us to more comprehensively examine differences in human behaviour between time-of-day and weekdays and weekends, and illustrates the importance of taking all these features into consideration for large-population studies. Non-surprisingly, our results suggest different wake-up times between weekdays and weekends; participants seem to wake up later during the weekends than weekdays. This information is of interest particularly given recent research suggesting that sleep irregularity may be a risk factor for cardio-metabolic disease (31). The large differences in movement and postural measures between weekdays and weekends suggest differences in the type of activities that participants partake in between weekdays and weekends. These differences are particularly striking when comparing women and men. We found that women spend more time with their wrist elevated above horizontal than men do (53% of their time vs. 36% for men). Similarly, the pitch and roll profiles coincide with increases in movement around noon of the weekend days, pointing towards a behavioural pattern that could be suggestive of “weekend warrior” lifestyle, where participants tend to do most of their physical activity during the weekend. Further inspection of the data through visualization techniques (figure 4 and associated video files) suggests that the activities participants engaged in strongly depended on time-of-day; it is apparent that the relative occupation of different physical spaces and the relationship between postures and movement changes drastically depending on the time of the day, indicative of engagement in different activity types.

We observed differences between men and women across most other substrata for both movement (vector magnitude) and posture (pitch and roll) measures, suggesting that men spent more time in postures that may be suggestive of sedentary behaviour than their female counterparts (sitting down, lying down). The inferred time estimate for sedentary behaviours (from vector magnitude and pitch), largely based on the methodology previously described by Rowlands et al (10), indicated that this was by far the most dominant behaviour across the whole population (∼17 hours/day). However, younger individuals tended to spend less time than their older counterparts in these sedentary behaviours (suggesting more active lifestyles), and even starker differences were observed between different BMI groups; individuals with higher BMI spent the most time in sedentary behaviours, and we statistically confirmed that this was independent of age and sex.

Movement and PAEE were both lower in the older age groups, a similar result to that observed in other population studies (2,32,33). We observed that older participants (60-65 age group) spend a large proportion of their time in postures that are similar to those with high BMIs, particularly in men. What was slightly paradoxical was that older and obese individuals spend more time at pitch angles generally associated with higher activity, ie with the arm below horizontal. As both movement and pitch are direct measurements of what the arm is physically doing, these results indicate true differences in activities, either as type or intensity or both. Using the sedentary time estimation methodology, it was suggested that older and heavier individuals spent more time in sedentary behaviours. Future inference work on raw non-dominant wrist acceleration signals may further elucidate other differences, for example in the specific type of activity performed, including the separation of awake sedentary behaviour and sleep.

Strengths of our study includes its standardised placement and 24-hour wear protocol which ensured greater certainty in the orientation of the accelerometer on each participant; that said, it is possible that some participants may have removed and replaced their device during the monitoring period. Still our results may provide guidance on probable axis orientation to other studies such as UK Biobank which do not have strict device orientation protocols. Another strength was that both wrist acceleration and PAEE was assessed simultaneously, thus providing more accurate stratification by PAEE levels; however a limitation of our work is that we only measured physical activity during one week of monitoring, and this may not be representative of habitual behaviour in this population. Another potential limitation is the separation between static and dynamic wrist acceleration; as has been previously addressed, the high-and low-pass filter parameters does not perfectly discriminate between static and dynamic and a small proportion of real movement will be missed during rapid rotations (34). Nonetheless, this is likely to only bias the movement differences we observe towards the null, since younger and slimmer individuals are more able to produce more rapid movements, and it will likely not impact much on the postural measures, as the gravitational acceleration component is several orders of magnitude larger than residual movement in the low-pass filtered signal, thus still returning a valid estimate of the relative distribution of gravity in the three axes.

### Conclusions

In conclusion, we found that direct measures of accelerometry-derived arm angles provide biomechanically meaningful information alongside the more well-established movement intensity metrics such as vector magnitude to better characterize objectively measured physical activity in free-living conditions. Movement is more likely to occur at arm angles below horizontal but despite older and heavier individuals moving less, these individuals still spend more time at lower arm angles, suggesting population differences in style of movement which may be important for other health outcomes.

### List of Abbreviations

PAEE: Physical Activity Energy Expenditure,
VM: Vector Magnitude,
VM HPF: Vector Magnitude High-Passed Filtered,
BMI: Body Max Index,
MET: Metabolic Equivalent Task

## DECLARATIONS

### Ethics approval and consent to participate

Ethics approval for the study was obtained from Cambridge University Human Biology Research Ethics Committee (Ref: HBREC/2015.16) with the ethical standards for human experimentation established by the Declaration of Helsinki. All participants provided written informed consent.

### Consent for publication

Not applicable

## Acknowledgements

We would like to thank the participants who took part in this study. We also thank the principal investigators of the Fenland study for their work on this population and the functional teams of the MRC Epidemiology Unit at Cambridge (Field Epidemiology, Study Coordination, Data management and IT) for supporting this study. The authors would like to thank Lewis Griffiths, Stefanie Hollidge and Antonia Smith for their assistance in the preparation of data for this study.

## Availability of data and materials

The data analysed during the current study are not publicly available because we have not obtained consent for public data sharing from the study participants; however data can be made available for analysis upon reasonable request to the corresponding author.

## Author’s contributions

IPP processed objective data, created the models, wrote the Python code, designed the analysis and wrote the manuscript. TW assisted on the analysis pipeline and provided technical advice on the design of the analysis and manuscript. KWe prepared data for use in this analysis. KWi provided advice on sedentary behaviours. NW and SB designed the study and obtained funding. SB assisted with the writing of the manuscript and provided technical advice on the analysis. All authors edited and approved the final version of the manuscript.

## APPENDIX

**Supplementary Figure 1:**
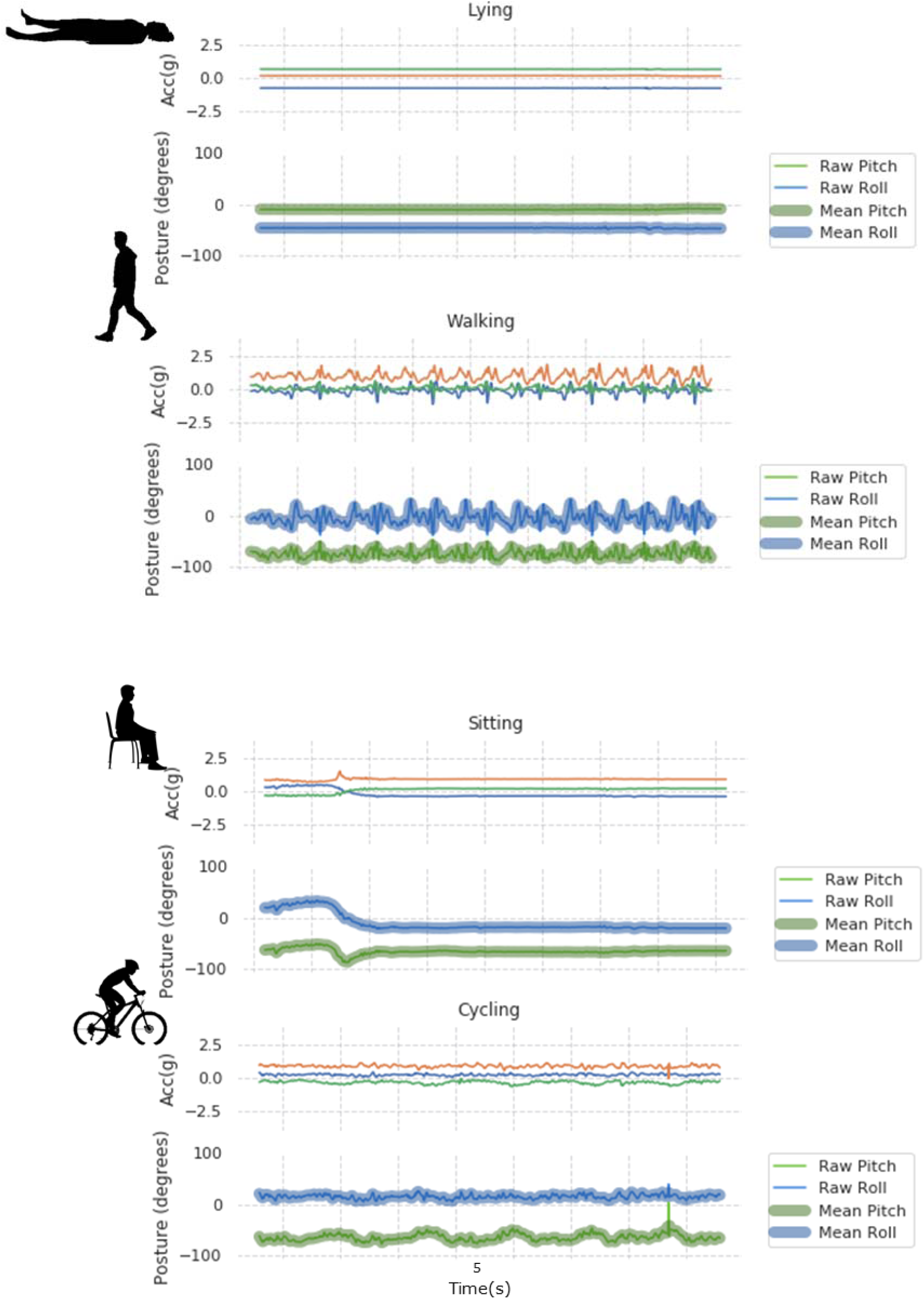
Raw triaxial wrist Acceleration, Pitch and Roll Profiles for typical daily activities. From top to bottom: lying, walking, sitting and cycling.

**Supplementary Figure 2:**
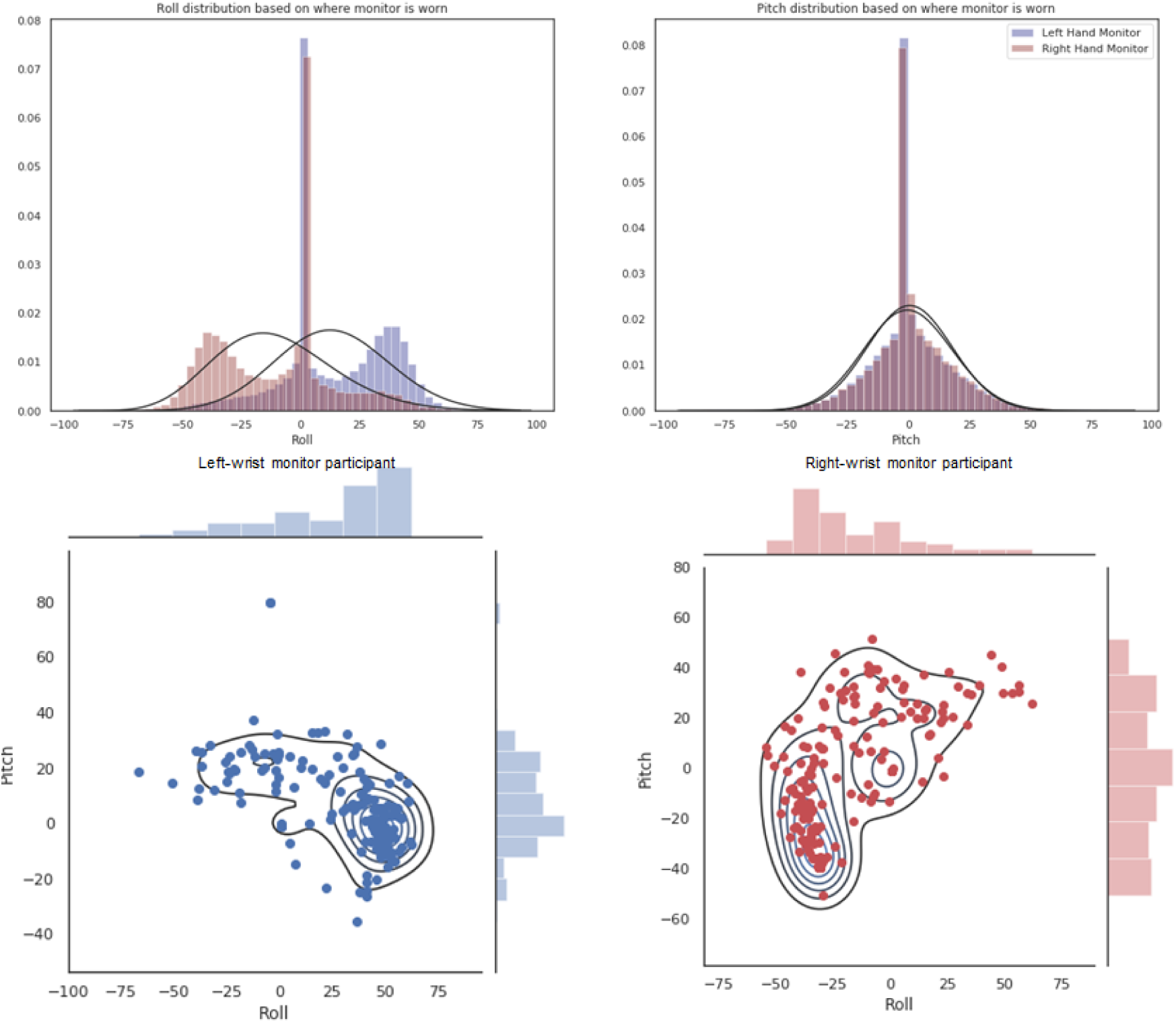
Untransformed Pitch and Roll distributions, stratified by left versus right-hand accelerometer wear (top panel). The two plots underneath show examples of pitch-roll distributions from participants wearing the accelerometer on their left (in blue) and right (in red) hand, respectively.

**Supplementary Figure 3:**
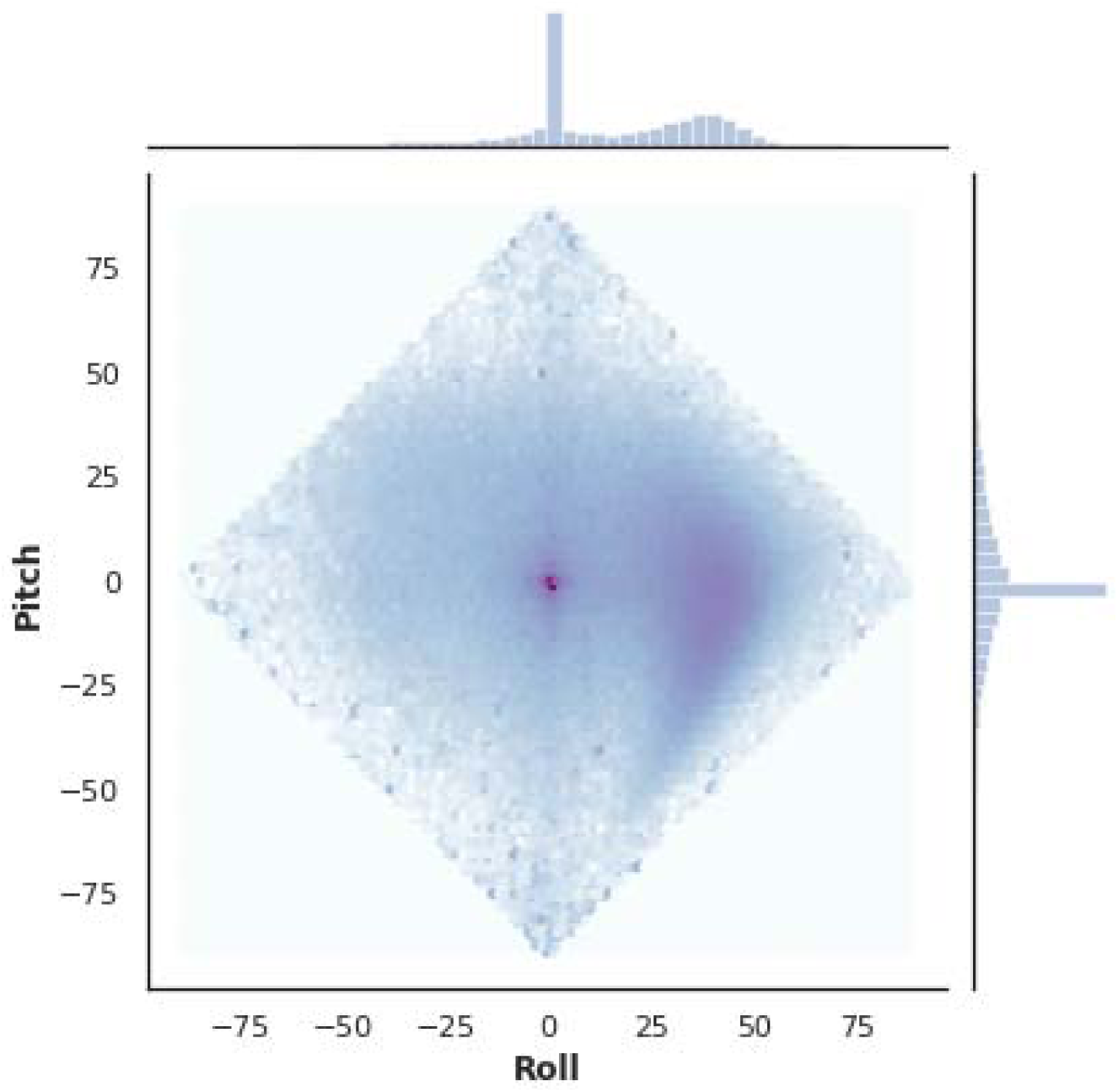
Two-dimensional distribution plot of Pitch and roll. Darker colours indicate higher occurrence of wrist positions

