## Supplementary figures and images for "Diurnal profiles of physical activity and postures derived from wrist-worn accelerometry in UK adults"

### Supplementary Figure 1

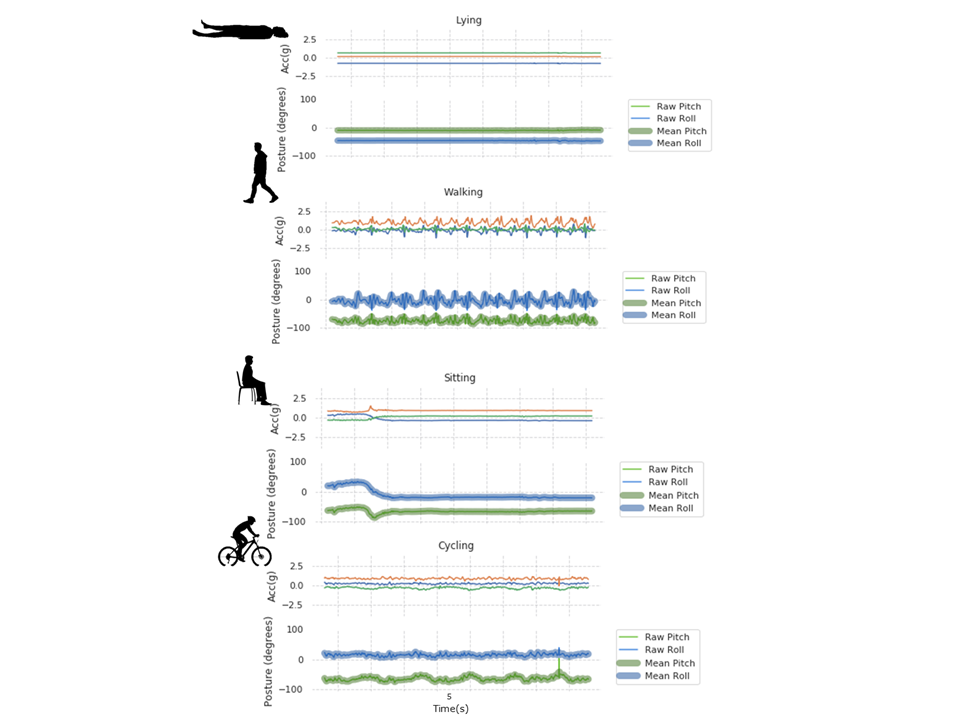

### Supplementary Material

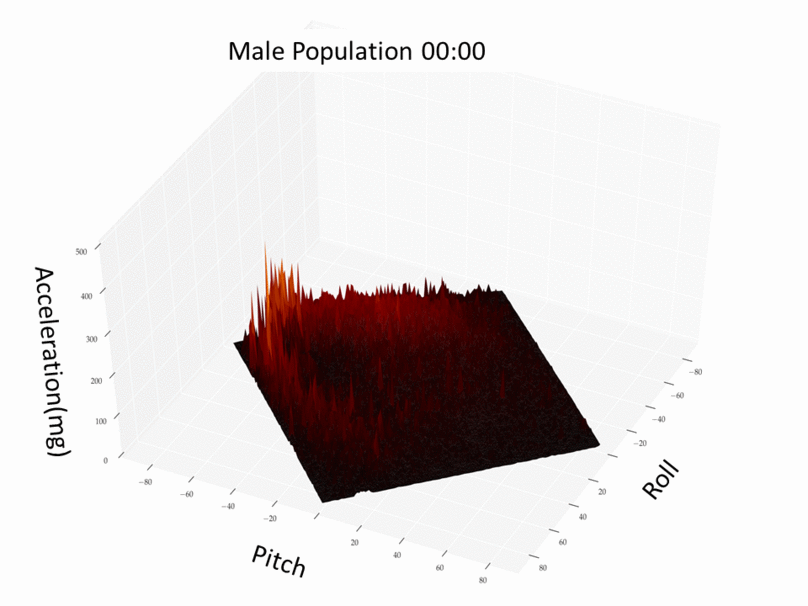

### Supplementary Material

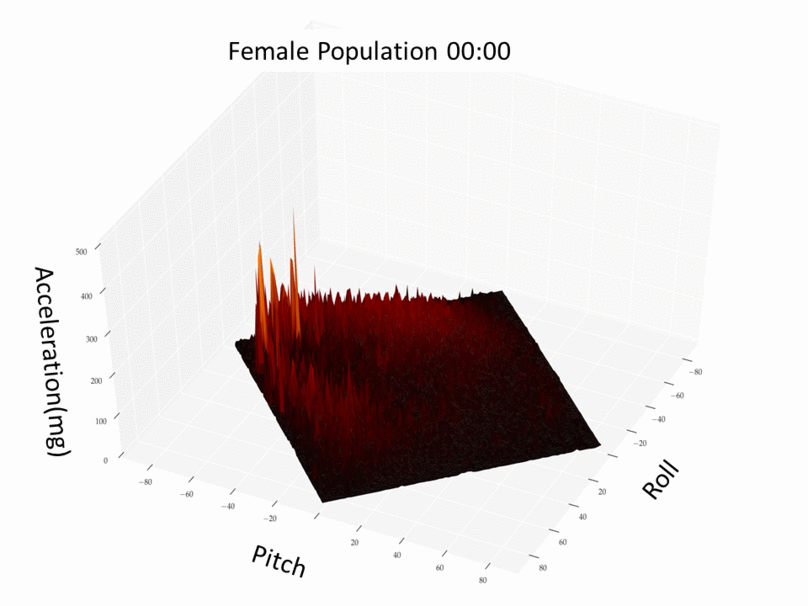
